# Predicting disordered regions driving phase separation of proteins under variable salt concentration

**DOI:** 10.1101/2022.11.26.518042

**Authors:** Esteban Meca, Anatol W. Fritsch, Juan M. Iglesias-Artola, Simone Reber, Barbara Wagner

## Abstract

We investigate intrinsically disordered regions (IDRs) of phase separating proteins regarding their impact on liquid-liquid phase separation (LLPS) of the full protein. Our theoretical approach uses a mean-field theory that accounts for sequence-dependent electrostatic interactions via a random-phase approximation (RPA) and in addition allows for variable salt concentration for the condensed and dilute protein phases. The numerical solution of the complete phase diagrams together with the tie lines that we derive for this model system leaves two parameters to be determined by fitting experimental data on concentrations of all species involved in the system. For our comparisons, we focus on two proteins, PGL-3 and FUS, known to undergo LLPS. For PGL-3 we predict that its long IDR near the C-terminus promotes LLPS, which we validate through direct comparison with *in vitro* experimental results under the same physiological conditions. For the structurally more complex protein FUS the role of the low complexity (LC) domain in LLPS has been intensively studied. Apart from the LC domain we here investigate theoretically two IDRs, one near the N-terminus and another near the C-terminus. Our theoretical analysis of these domains predict that the IDR at the N-terminus (aa 1-285) is the main driver of LLPS of FUS by comparison to *in vitro* experiments of the full length protein under the same physiological temperature and salt conditions.

**SIGNIFICANCE:** Intrinsically disordered proteins are drivers of cellular liquid-liquid phase separation. However, it remains a challenge to directly predict the phase behaviour of a protein based on its primary sequence, and under physiological conditions. We present a random-phase approximation that allows for variable salt concentration and thus accounts for salt partitioning. We use this to link the sequence of the disordered regions with the behaviour of the complete protein through direct comparisons to *in vitro* phase-separation assays. In particular, for FUS we determine the exact region responsible for LLPS, weighting in a long-standing debate.

## INTRODUCTION

Protein condensation driven by liquid-liquid phase separation (LLPS) is a powerful concept to understand the mesoscale organization of cells. It provides a simple mechanism to form non-membrane bound organelles that separate from the nucleo- and cytosol. Examples of such biomolecular condensates include nucleoli, P-granules, stress granules and centrosomes as reviewed in Banani et al. (1), Shin and Brangwynne (2), Alberti and Hyman (3). These condensates correspond to a protein-rich phase coexisting with a protein-poor bulk phase.

One of the drivers of cellular LLPS are intrinsically disordered regions (IDRs), these can be either part of a protein or constitute the entire protein (intrinsically disordered proteins, IDPs). IDRs are highly dynamic regions within protein sequences that lack stable secondary or tertiary structure. Yet, they facilitate weak multivalent interactions. On the sequence level, driving forces include electrostatic interactions between charged motifs that promote long- and short-range interactions. Short-range interactions are characterized by directional interactions of dipoles or positive charges with aromatic groups (4). Thus, the phase behaviour of a given protein is sequence encoded. Condensation, however, does not only depend on protein structure but also on the environmental conditions. These include temperature, ionic strength of the constituents or their concentration. For example, condensates can dissolve upon raising temperature or salt concentration and can reform when conditions are reverted.

Despite increasing experimental evidence, lending insight into the physical chemistry that is driving LLPS, it remains a challenge to directly predict the phase behaviour of a protein based on its primary sequence and solvent environment. This limits our ability to predict how changes in the amino acid sequence of a protein influence its phase behaviour. Therefore, theoretical predictions for phase diagrams are needed to guide experimental research and provide insights into the molecular basis of physiological and pathological processes related to diseases and ageing.

While molecular dynamics simulations provide detailed biophysical information on a single protein level (5), they quickly become computationally expensive when applied to large ensembles of phase-separating proteins in a solvent. A coarse-grained lattice-based approach is the classical Flory-Huggins theory (6, 7) and its extension to the Voorn-Overbeek theory (8), which incorporates electrostatic interactions via the Debye-Hückel theory. In the derivation of these theories variations in charge patterns that are responsible for phase-separation are averaged out. They thus cannot capture the structure- or even sequence-specific phase behaviour, which is a signature of IDPs. On an intermediate coarse-grained scale, Field Theoretic Simulations (FTS), rooted in statistical mechanics, are able to incorporate structural information of proteins and has recently been used to predict LLPS of tau, see e.g. Zhang et al. (9). However, since FTS relies on the full partition function of the free energy it is still numerically very demanding. Asymptotic approximations of the full partition function, such as the Random-Phase Approximation (RPA), reliably account for the structural features of proteins. Indeed, Lin et al. (10, 11) have pioneered the application of RPA to LLPS of phase separating proteins to predict the sequence-specific phase behaviour of the RNA helicase Ddx4, see also the recent review by Dinic et al. (12).

Here we predict the phase behaviour of proteins by using a thermodynamically consistent theoretical mean-field model that includes salt concentration as a variable. Using a random-phase approximation, we introduce the sequence-dependent electrostatic interactions arising only from one IDR at a time to build the free energy. In our analysis, we assume that this free energy encodes the phase behaviour of the complete protein, and thus derive temperature-protein concentration and salt-protein concentration phase diagrams. Under the same physiological conditions we directly compare to *in vitro* experiments. We study in depth two well-known proteins on which this approach is successful. These are PGL-3 (*C. elegans*) and fused in sarcoma (FUS). These two proteins are known to undergo LLPS *in vitro* and *in vivo* (13, 14). By matching to the exact conditions of *in vitro* experiments, we determine the IDRs that drive LLPS of these proteins.

We first investigate PGL-3, since its phase behaviour is well understood (Brangwynne et al. (13), Saha et al. (15)) and validate our predictions with experimental data of the dilute and condensed phase concentration from *in vitro* studies under physiological salt conditions. Our results confirm that the IDR at the C-terminus drives LLPS of PGL-3. We then focus our analysis on FUS, where the driving forces and sequence domains responsible for LLPS are still under debate (Wang et al. (14), Patel et al. (16)). We identify and analyse three regions as possible candidates to impact the phase behaviour of FUS, the LC region, an IDR at the N-terminus and an IDR at the C-terminus.

Our analysis reveal that the domain at the N-terminus, from amino acids 1 to 285, to be responsible for LLPS when comparing to *in vitro* experimental data of FUS.

## MATERIALS AND METHODS

### PGL-3 and PGL3-GFP protein purification

PGL-3 was purified from insect cells according to (15). SF9-ESF cells were infected with baculovirus containing the PGL-3-GFP-6HIS protein under the polyhedrin promoter. Cells were harvested after 3 days of infection by centrifugation at 500 x g for 10 min and then resuspended in lysis buffer (25 mM HEPES 7.25, 300 mM KCl, 10 mM imidazole, 1 mM DTT, 1 protease inhibitor). Cells were lysed by passing the cells 2 times through the LM20 microfluidizer at 15 000 psi. The lysate was then centrifuged at 20 000 rpm for 45 min at 15 °C. The lysate was loaded in a pre-equilibrated Ni-NTA column with lysis buffer at 3 mL/min. The Ni-NTA column was rinsed with 10 C.V of wash buffer (25 mM HEPES 7.25, 300 mM KCl, 20 mM imidazole, 1 mM DTT, 1) and the protein was eluted in 1.5 mL fractions with elution buffer (25 mM HEPES 7.25, 300 mM KCl, 250 mM imidazole, 1 mM DTT). After elution the GFP tagged was cleaved to produce untagged PGL-3. The cleavage was performed using a TEV protease overnight at 4 °C. PGL-3 and PGL-3-GFP proteins were diluted with Dilution buffer (25 mM Tris pH 8.0, 1 mM DTT) to reach 50 mM KCl before loading the protein in an anion exchange HiTrapQ HP 5 mL column. The HiTrap column was previously equilibrated first with HiTrapQ elution buffer (25 mM Tris pH 8.0, 50 mM KCl, 1 mM DTT) and then with HiTrapQ binding buffer (25 mM Tris pH 8.0, 1 M KCl, 1 mM DTT). The column was mounted in a Äkta Pure FPLC system. After the sample was loaded the column was washed with HiTrapQ binding buffer. The sample was finally eluted with a linear gradient from 0 to 55% of HiTrapQ elution buffer (25 mM Tris pH 8.0, 1 M KCl 1 mM DTT) for 25 C.V. Finally a 100% HiTrap elution buffer step was performed for 5 C.V. The pooled fractions were then loaded in a HiLoad 16/60 Superdex 200 size exclusion chromatography column that was previously equilibrated with superdex buffer (25 mM HEPES 7.25, 300 mM KCl, 1 mM DTT). After size exclusion, the final samples were collected.

### FUS protein purification

Unlabeled FUS purified from a baculovirus construct containing N-HIS-MBP-FUS-TEV-SNAP. SF9-ESF cells were harvested after three days of infection by centrifugation at 500 x g for 10 min. The cell pellet was resuspended using 50 mL of lysis buffer (50 mM Tris pH 7.4, 500 mM KCl, 5% glycerol, 10 mM imidazole, 1mM PMSF, 1X protease inhibitor) for every 50 mL of cultured cells. The cells were lysed by passing them 2 times through the LM20 microfluidizer at 15 000 psi. The lysate was then centrifuged at 20 000 rpm for 45 min at 15 °C. The supernatant was collected and loaded into a Ni-NTA column that was previously equilibrated with lysis buffer. After loading the sample the column was washed for 10 C.V. with Ni-NTA was buffer (50 mM Tris pH 7.4, 500 mM KCl, 5% glycerol, 20 mM imidazole). The protein was then eluted with Ni-NTA elution buffer (50 mM Tris pH 7.4, 500 mM KCl, 5% glycerol, 300 mM imidazole). The collected fractions where then loaded into a MBPTrap HP column preequilibrated with Ni-NTA elution buffer. The MBP column was washed for 10 C.V with MBP wash buffer (50 mM Tris pH 7.4, 500 mM KCl, 5% glycerol). After washing, the sample was eluted with MBP elution buffer (50 mM Tris pH 7.4, 500 mM KCl, 5% glycerol, 500 mM arginine, 20 mM maltose). The protein was diluted to a concentration of less than 15 *μ*M using MBP elution buffer. 3C and TEV proteases were then added to cleave the MBP and SNAP tags from the FUS construct. The cleavage reactions were incubated overnight at 18 °C. Finally the protein was loaded in a SepFast GF-HS-L 26 × 600 mm gel filtration to remove the cleaved MBP and SNAP tags and exchange the buffer. The SepFast column was previously equilibrated in storage buffer (50 mM HEPES pH 7.25, 750 mM KCl, 5% glycerol, 1 mM DTT). The sample was concentrated to a final concentration of 15 *μ*L using 30 kDa Amicon centrifuge filters. FUS-GFP was purified as previously described in (14).

In our analysis we mainly use the MetaDisorder predictor by Kozlowski and Bujnicki (17) to identify the disordered and low complexity regions. MetaDisorder queries other predictors and generates a consensus answer, with an algorithm that tests the strength of each method against several datasets. It thus addresses the issue of training-set dependent model predictions. In particular, MetaDisorder includes also widely used IUpred predictors (18).

### Measurement of *c*_*out*_ and *c*_*in*_

A master-mix of 95% unlabeled and 5% labeled protein was prepared from high-salt stock solutions (PGL-3: 300mM KCl, 25mM HEPES, 1mM DTT; FUS: 750mM KCl, 25mM HEPES, 1mM DTT, 5% Glycerol). Phase separation was initiated by diluting the stock salt concentration directly before encapsulation in water-in-oil emulsions that were created using Pico-Surf (2% (w/w) in Novec 7500, Sphere Fluidics). Emulsions were loaded on a temperature controlled stage and the sample chambers were sealed with a two-component silicone (Picodent, Twinsil Speed). After a 30 min waiting time at the desired temperature the emulsion droplets were imaged using a 40X, 0.95 N.A., air objective mounted on an Olympus IX83 microscope stand controlled via CellSens. Confocal Z-stacks were recorded using a Hamamatsu Orca Flash 4.0 connected to a Yokogawa W1 spinning disc unit. Large 3D tile-images were collected to increase the statistics of individual emulsion droplets.

The fluorescence and bright-field images were analyzed using a custom MATLAB code. This allowed us to derive the volume fraction *V* _*frac*_ of the condensed phase in each emulsion droplet by image segmentation. Using volume and mass conservation we can derive a linear relationship volume fraction and total protein concentration *c*_*tot*_ : *V* _*f rac*_ = 1 /(*c*_*in*_ −*c*_*out*_) · *c*_*tot*_ −*c*_*out*_ /(*c*_*in*_ −*c*_*out*_). We then used a set of total protein concentrations to determine both *c*_*out*_ and *c*_*in*_ via linear regression to this equation. Thus, this allows for experimental measurements of temperature and salt dependent phase diagrams using small amounts of protein sample.

### Field theoretic approach

For the purpose of this study, i.e. analysing the impact of different domains of an IDP on its propensity to phase separate, we use field theoretic approaches rooted in statistical mechanics that are able to incorporate detailed structural information of polyampholytes such as proteins. They are obtained from the partition function for ensembles of coarse-grained polyampholytes, which are represented, via the Hubbard-Stratonovich transformation, as multi-dimensional functional integral (path integral) over all possible states or the polymeric system. Field theoretic simulations (FTS) are numerical methods that consider the full functional integral of the associated partition function. However, the integration of the full integral relies on stochastic sampling via Monte-Carlo methods that have shown considerable convergence problems due to the oscillatory nature of the resulting distribution function (19). Nevertheless, this approach has recently been used successfully for the tau protein (9). It holds promise for the development of an appropriate and efficient theory that will allow to characterize a whole class of proteins such as IDPs and proteins that contain intrinsically disordered domains, and can be used to deliver analytical insight into the underlying biophysical principles leading to LLPS. For our analysis we use the Random-Phase Approximation (RPA). As is the case for self-consistent field theory, it can be derived from a saddle-point approximation, taking into account the asymptotically dominant contribution (a single Gaussian distribution) of the functional integral of the partition function, reducing the field theory to a mean-field model where a single Gaussian configuration interacts with an average effective field. It is one of the simplest analytical theories, that can account for small site-specific fluctuations, e.g. of charge patterns and structural features of the protein.

The RPA approach we use here has been set-up previously by Lin et al. (10, 20) for the IDR of Ddx4. For our analysis we use the extended model that includes salt as an additional variable, apart from the protein (here for IDRs of PGL-3 and FUS) (*ϕ*_*aa*_), counterions (*ϕ*_*c*_), KCl (*ϕ*_*s*_) and water. The free energy

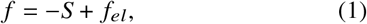

combines the entropic part *S*, coming from Flory-Huggins solution theory:

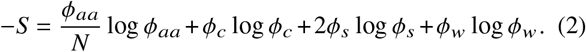

where *N* denotes the number of amino acids and the solvent fraction can be written as *ϕ*_*w*_ = 1 − *ϕ*_*aa*_ − *ϕ*_*c*_ − 2*ϕ*_*s*_, with the electrostatic part of the free energy *f*_*el*_. Note that we only have a single volume fraction *ϕ*_*s*_ to represent the salt concentration, which implies that the concentration of both ions conforming the salt is considered equal in all phases. Here we follow Lin et al. (10), but this assumption is not necessarily correct, see the discussion.

For the evaluation of *f*_*el*_ the Random Phase Approximation is commonly used, (21–24). It can be expressed in its simplest non-dimensional form as the following integral:

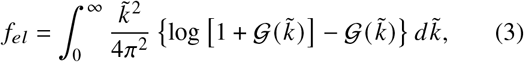

with

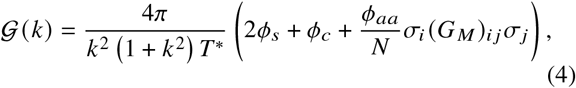

where *T* ^*^ is the nondimensionalised temperature which includes the relative permittivity of the medium. The correlation matrix for the chain can be written as

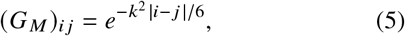

where the lengths have been nondimensionalized with the characteristic length *a* of the polymeric link of the associated Gaussian chain, here the domain of interest for protein PGL-3 or FUS. Note, that the neutrality condition implies *ϕ*_*c*_ = |*σ*|*ϕ*_*aa*_, where 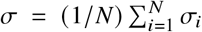 is the average charge density of the protein, and *σ*_*i*_ is the charge of each of its amino acids. Together, this reduces the system to two quantities to be solved for, *ϕ*_*aa*_ and *ϕ*_*s*_.

### Computation of the phase diagrams

To compute the phase diagrams including salt dependence, in particular when solving for the tie lines of the model system, we integrate the system using a Gauss-Laguerre quadrature, with the points computed using both our own implementation of the classic method by (25) and the state-of-the-art method by (26). In order to find the tie lines between two coexisting points (*ϕ*_*aa*_, *ϕ*_*s*_)|_*α*_ and (*ϕ*_*aa*_, *ϕ*_*s*_)|_*β*_, we solve the following system, which is equivalent to the equality of the electrochemical potentials and the common tangent construction:

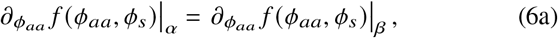

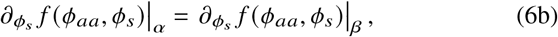

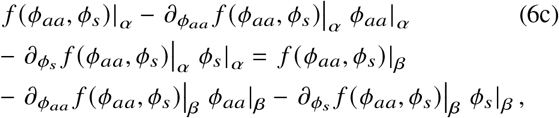

where the nonlinear system of equations is solved by using the *trust-region-dogleg* algorithm as implemented in Octave.

Note that this is a system of three equations with four unknowns, the volume fraction of salt and protein in the dilute and in the condensed phases. Because of the way in which the protein concentrations are measured experimentally, we know the salt concentration in the dilute phase, which allows the system of equations to be determined when comparing with the experiments.

In order to draw the complete phase diagram, we can take one of the unknowns as a parameter and solve for the others. In order to solve effectively the system above it is imperative to develop a continuation strategy. Naively, one could vary the unknown taken as a parameter and solve for the others using the prior solution as a guess. This strategy is known as *natural parameter continuation* (27), but the system above presents turning points for the parameter, which requires the development of a *pseudo-arclength continuation* algorithm. The latter implies the definition of a new variable, the pseudo arclength, which is defined as the arclength of the solution curve in the four-dimensional space spanned by the four unknowns in the system above. In practical terms, using this method implies adding an additional equation to the previous system that imposes a fixed increase of the arclength from the prior solution. Thus, we obtain a system of four solutions with four unknowns, that is similarly solved using the *trust-region-dogleg* algorithm.

### Parameter estimation and fitting procedure

We now express T* and *ϕ*_*aa*_ in terms of experimentally accessible variables, concentrations (*c*) and temperatures (*T*). In the case of *ϕ*_*aa*_, we know that it corresponds to the total volume occupied by amino acids over the volume of the total lattice. We take the volume of a lattice site to be that of a single water molecule, which corresponds to 1/55.5M. We would have then for PGL-3 in the presence of KCl (note that PGL-3 consists of 693 amino acids):

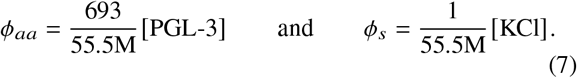

There are two implicit assumptions in this calculation. One, it is assumed that each amino acid takes the same volume as a water molecule, which is clearly not true, and two, that the volume of a protein scales linearly with the number of amino acids. The latter can be argued to be false, since different configurations of the protein will give different volumes. Therefore, there is not a clear and easy way of relating the volume fraction and the protein concentration.

The non-dimensional temperature *T* ^*^ is related to *T* by

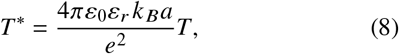

where *ε*_0_ is the vacuum permittivity, *ε*_*r*_ is the relative permittivity of the medium, *k* _*B*_ is Boltzmann’s constant, *a* the link length and *e* is the charge of the electron. The two unknowns in Eq. (8) are *a* and *ϵ*_*r*_. We take *a* to be the C*α*-C*α* virtual bond length of 3.8Å and fit *ε*_*r*_, due to the lack of a complete theory to derive its value, we only know that it should be between 2 (typical value for hydrocarbon crystals) and 80 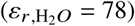.

The impossibility of obtaining accurate theoretical estimates for the relation between volume fraction and the concentration of the protein and for the permittivity of the solution make it necessary to proceed with a fitting procedure. There exist two parameters, *β*_1_ and *β*_2_ that allow us to scale the volume fraction and the non-dimensional temperature to fit the data of a *T* -*c* phase diagram, i.e. *T* = *β*_1_*T* ^*^ and *c* = *β*_2_*ϕ*_*aa*_. Once the phase diagram is computed, we obtain a functional relation between *T* ^*^ and *ϕ*_*aa*_, *ϕ*_*aa*_ = *f* (*T*^*^) (note that *f* has different branches, a dilute and a condensed branch). If we know both parameters we can rescale the previous relation to obtain *c* = *β*_2_ *f* (*T*/*β*_1_), a predictor for the concentration. We can find both parameters by minimizing the following function:

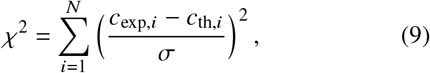

where *c*_exp_ are the experimental values of the concentration, *σ* is the standard deviation of the experimental points. The sum runs over all experimental points including all branches and all values of the salt concentration. The parameter *β*_1_ is allowed to vary with the salt concentration on account of the strong dependence of the permittivity on salt concentration. Note that we do not assume any particular functional dependencence of *ε*_*r*_ on salt, we simply fit a value of *β*_1_ at each salt concentration. On the other hand, *β*_2_ is considered independent of salt concentration, which is the most parsimonious choice. Finally, note that the lack of a good theory for the dependencence of *ε*_*r*_ on salt implies that a single value of *β*_1_ has to be selected for the computation of the salt-concentration phase diagrams, which limits the agreement of the model and the experimental data.

Eq. (9) is minimized using a sequential quadratic programming algorithm as implemented in Octave. The parameter *χ*^2^ will provide then a measure of the goodness of fit for each case. In Table 1 we give a summary of all fitting parameters for all cases we have considered.

**Table 1:** Fitting parameters for the IDRs of FUS and PGL-3 at KCl and temperatures used in experiments. The temperature *T* is scaled via the parameter *β*_1_ (K) and the concentration *c* is scaled via *β*_2_ (mM).

| IDR | [KCl]<br>(mM) | T<br>$\beta_1$ (K) | c<br>$\beta_2$ (mM) | $\chi^2$ | $\epsilon_r$ |
| --- | --- | --- | --- | --- | --- |
| FUS-C1 | 100 | $5.35 \times 10^3$ | 10.1 | 191 | 8.22 |
| | 150 | $5.77 \times 10^3$ | 10.1 | 191 | 7.62 |
| | 200 | $6.01 \times 10^3$ | 10.1 | 191 | 7.31 |
| FUS-N1 | 100 | $4.48 \times 10^3$ | 207 | 3.35 | 9.82 |
| | 150 | $5.02 \times 10^3$ | 207 | 3.35 | 8.76 |
| | 200 | $5.36 \times 10^3$ | 207 | 3.35 | 8.21 |
| FUS-LC* | 100 | $7.90 \times 10^4$ | 25.3 | 3.92 | 0.556 |
| | 150 | $1.10 \times 10^5$ | 25.3 | 3.92 | 0.400 |
| | 200 | $1.38 \times 10^5$ | 25.3 | 3.92 | 0.317 |
| PGL-3-C1 | 100 | $1.15 \times 10^3$ | 16.2 | 23.4 | 38.2 |
| | 150 | $1.31 \times 10^3$ | 16.2 | 23.4 | 33.6 |
| | 200 | $1.35 \times 10^3$ | 16.2 | 23.4 | 32.6 |

\* We note that the parameter values for FUS-LC correspond to the constant salinity case, since there is no binodal in the partitioning case for FUS-LC. For any KCl used, the relative permittivity $\epsilon_r$ is smaller than 1.

## RESULTS

The phase behavior of a given protein is encoded in its free energy function *f*,

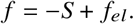

As indicated in Fig. 1 A, the free energy has an entropic part *S*, representing the Flory-Huggins interactions

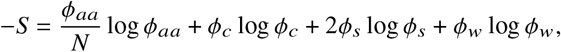

where *N* denotes the number of amino acids. Apart from the volume fraction of the amino acids of the protein *ϕ*_*aa*_, the volume fraction of counterions *ϕ*_*c*_ and the volume fraction *ϕ*_*w*_ of water, we also include in our analysis the volume fraction of salt ions KCl (*ϕ*_*s*_) as an additional variable. The second term *f*_*el*_ represents the electrostatic multivalent interactions of protein chains with each other and the surrounding salt solution, and in our case drives LLPS. Thus, it is this part of the free energy for which the Random Phase Approximation (RPA) is being applied in order to account for the dependence of the free energy on the protein structure (see Field theoretic approach in the Methods and Materials section). After we determine the disorder tendency of the structure of the protein structure, we derive the temperature-protein concentration and salt-protein concentration phase diagrams, as sketched in Fig. 1 B, based on the free energy function *f* for the proteins PGL-3 and FUS (see Computation of phase diagrams in Methods and Materials section).

**Figure 1:**
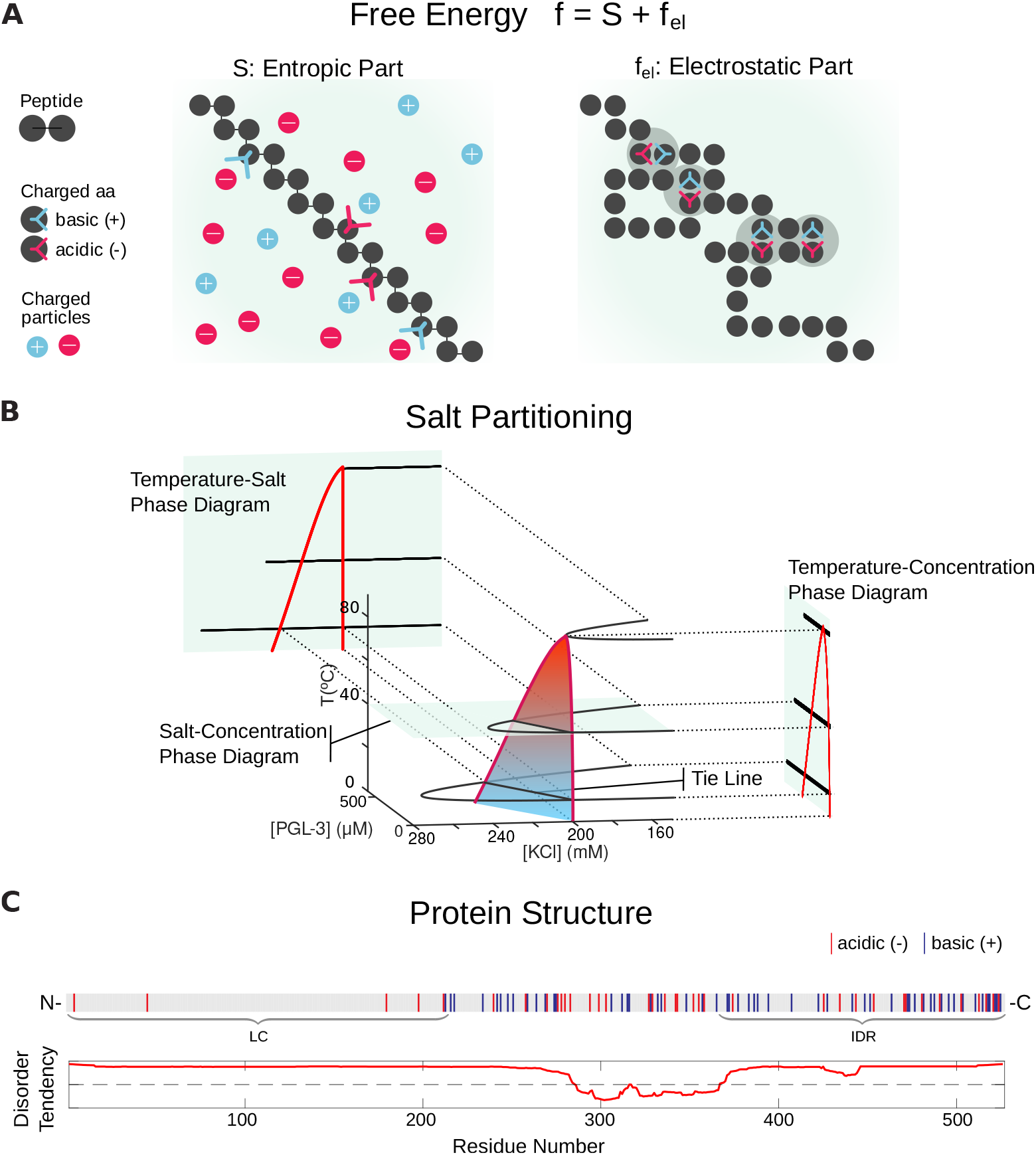
Theoretical approach **A**: We assume short range entropic and long range electrostatic interactions with residues of a peptide chain taking part in multivalent interactions, combine to drive liquid-liquid phase separation in solution. **B**: Random Phase Approximation (RPA) we present allows for variable salt concentration and can account for salt partitioning. The corresponding phase diagrams typically show *pointy* shapes as a consequence of a discontinuity in the difference in salt between the condensed and dilute phases at high temperatures. **C**: For the sequence analysis we use the MetaDisorder predictor MD2 by Kozlowski and Bujnicki (17) to identify the disordered and low complexity regions (represented here is FUS). MetaDisorder queries other predictors and generates a consensus answer, with an algorithm that tests the strength of each method against several datasets. It thus addresses the issue of training-set dependent model predictions. In particular, it also includes other widely used predictors such as IUpred (18).

### LLPS in PGL-3 is accompanied by salt partitioning

We first test the predictions of this model with experimental results for a range of different bulk concentrations of recombinant PGL-3 in an *in vitro* phase-separation assay. The experiments exhibit the generic behaviour, that higher PGL-3 concentrations are necessary at increasing temperatures in order to initiate LLPS at physiological salt (150 mM KCl) concentration. This is quantified at each temperature with a corresponding protein saturation concentration *c*_*out*_ (Fig. 2 B,C).

**Figure 2:**
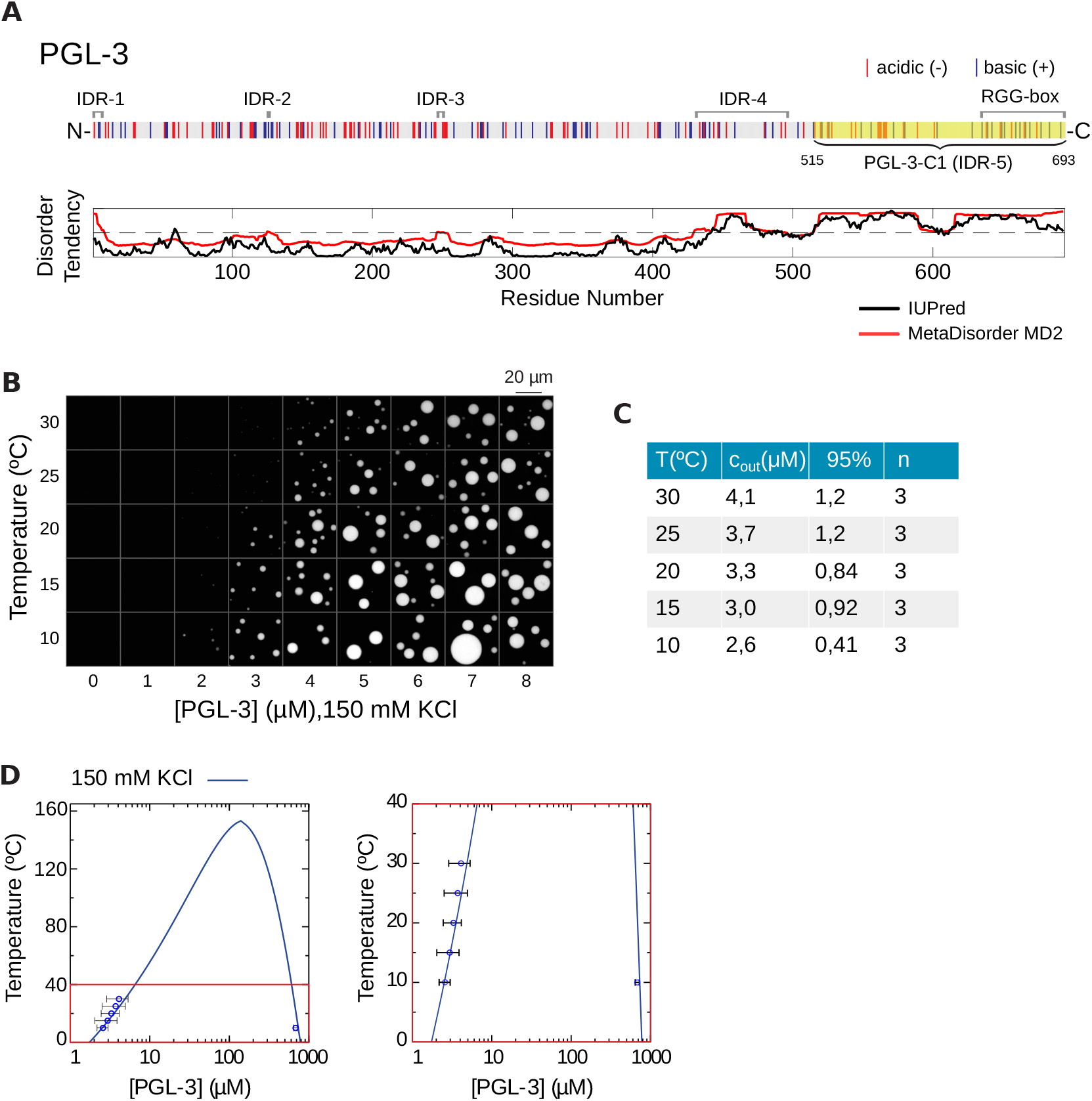
Phase separation of PGL-3 **A** Sequence of *C. elegans* PGL-3 with negatively (red) and positively (blue) charged amino acids. MetaDisorder MD2 (red) and IUPred (black) identify five disordered regions with IDR-5 (PGL-3-C1) at the C-terminus being the longest and RGG-box as in (15). PGL-3-C1 (highlighted) is used for the computations of the phase diagrams below. **B** Experimental data show the temperature- and concentration-dependent phase separation of PGL-3 at physiological salt conditions of 150 mM KCl. **C** Quantification of data in **B** to derive *c*_*out*_. *n* is the number of repetitions of the experiment, 95% corresponds to the confidence interval. **D** Predicted temperature-protein concentration phase diagram for PGL-3-C1 based on RPA. The phase diagram is computed with *ε*_*r*_ = 33.6 and *β*_2_=16.2 mM, using Eqs. (2)-(7) (see Methods and Materials). To the right is a zoom of the red region.

To investigate the role of intrinsically disordered regions (IDRs) of PGL-3, we determine (using MetaDisorder MD2) four small disordered regions (IDRs) and one large IDR (IDR-5: aa 515-693) near the C-terminus, which we denote by PGL-3-C1, see Fig. 2 A. It is the longest linear sequence predicted to lack a secondary structure. Indeed, this region has previously been shown to include a set of six C-terminal RGG repeats (aa 633-695), which bind RNA and promote droplet formation Saha et al. (15). For the derivation of the binodal, specifically the RPA free energy due to the electrostatic interactions (*f*_*el*_), we use the longest IDR PGL-3-C1.

For the quantitative comparison, we need to relate the PGL-3 volume fraction *ϕ*_*aa*_ and the non-dimensional temperature *T* ^*^ in terms of the experimentally accessible PGL-3 concentration (*c*) and temperature (*T*), respectively.

To relate *ϕ*_*aa*_ with *c* we could assume, as it is sometimes done, that we have a lattice with each lattice site having the volume of a water molecule and that an amino acid occupies exactly one site. This provides a conversion factor (Eq. 7), but it is at best a crude estimation. Similarly, the non-dimensional temperature depends on the relative permittivity *ε*_*r*_ (see Eq. 8). A complete theory to derive the value of *ε*_*r*_ is lacking, thus leaving is with little choice but to fit both conversion factors *β*_1_ and *β*_2_ for *T* = *β*_1_*T* ^*^ and *c* = *β*_2_*ϕ*_*aa*_. Note that *ε*_*r*_ is expected to have a strong dependence on salt concentration, and hence we will fit *β*_1_ independently for each temperature phase diagram at each salinity. (See Materials and Methods for the details).

The experimental values were obtained using a method that calculates *c*_*in*_ and *c*_*out*_ by measuring the volume of the condensed phase in an enclosed compartment with fluorescence microscopy. This method relies on the linear relationship between total protein concentration and condensed phase volume fraction. A visual example of this linear relationship can be observed in Fig. 2 B for PGL-3 at different temperatures. Fig. 2 B shows that an increase in temperature is accompanied by an increase in the proetin concentration required for phase separation (*c*_*out*_). Here, we used water-in-oil emulsions to encapsulate the protein solutions immediately after triggering phase separation. The corresponding experimental values of *c*_*out*_ are presented in the table in Fig. 2 C.

The theoretical phase diagram in Fig. 2 D reaches its maximum near 160°C. At this point the slope of the phase diagram is clearly discontinuous, thus creating a *pointy* feature. In the zoom on the temperature range of the experimental data (Fig 2 D, right panel), we observe a good agreement between theory and experimental data, particularly for lower temperatures. The value of the concentration for the theoretical condensed branch at 10°C does not fall strictly within the 95% confidence interval, but it is nevertheless close. Considering that we only have two parameters at our disposal, we can consider this agreement of experiments and theory very good. The goodness of fit is quantified by the comparatively low value of the *χ*^2^ parameter (see Table 1 in Methods and Materials).

Our theoretical model allows for a variable salt concentration. As a consequence, the protein-poor and protein-rich phases may have different equilibrium salt concentrations, which implies in turn that the tie lines connecting both equilibria have a non-zero slope. We note that this is responsible for the characteristic *pointy* shape of the temperature phase diagram in Fig. 2 D, which is a path along the complete 3D phase diagram (shown in the supplemental Fig. S1). Experimentally, the value of the saturation concentration *c*_*out*_ is found by extrapolation at a value where the salt concentration in the supernatant phase is fixed. This implies a constant salt concentration in the protein-poor branch but a varying salt concentration in the protein-rich branch. This is the underlying cause of the non-smoothness of the binodal curve at the point where both branches meet.

Starting with the protein-poor branch we measure how salt affects *c*_*out*_. Again we obtain a generic behaviour where higher salt concentration requires higher protein concentration for phase separation of PGL-3 to occur (Fig. 3 A). Compared to the dependence we saw for temperature, however, our experimental results suggest that salt has a stronger influence on *c*_*out*_. Specifically, in the range from 100 to 220 mM KCl at 20 °C, *c*_*out*_ changes approximately 30-fold from 1.12 to 34 *μ*M (Fig. 3 B). For the quantitative comparison against experimental results for the salt-concentration phase diagram we use the now already determined scaling parameters to derive the phase diagram from our theoretical model. For that purpose we use the scaling parameters fitted at a constant salt concentration of 150 mM KCl, since a model that captures how permittivity changes with salt concentration is not available. The resulting theoretical curve at 20°C (Fig. 3 C) has a maximum near 250 mM KCl, a salinity above which we do not expect LLPS The agreement of the overlaid experimental points with the theoretical dilute branch (Fig. 3 C, right panel) is very good, with the discrepancy with the confidence interval stemming from the fact that the value of the permittivity varies strongly with salt concentration, and we are considering it to be constant. At the lower temperature of 10°C we observe a similar good agreement between theory and experiments in the dilute branch (Fig. 3 D) and on the condensed branch, again considering we only have two parameters at our disposal.

**Figure 3:**
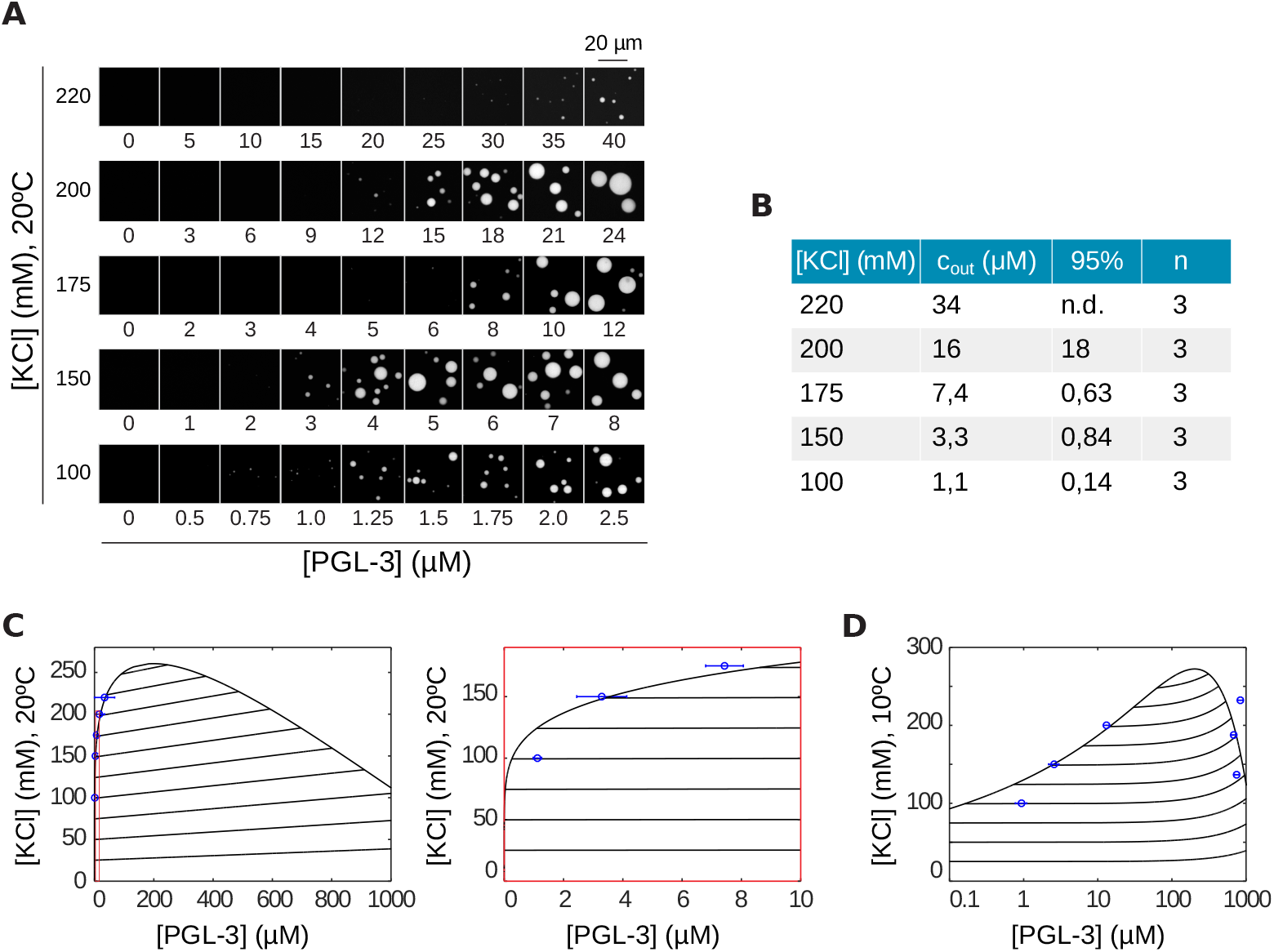
Salt partitioning and the domain PGL-3-C1 capture LLPS for PGL-3. **A** Salt- and concentration-dependent phase separation of PGL-3 at 20 °C. They show higher concentration of PGL-3 is needed for higher salt concentrations in order to initiate phase separation. **B** Quantification of data in **A** to derive *c*_*out*_ with *n* = 3 repetitions, 95% denotes the corresponding confidence interval. **C** Predicted salt-concentration phase diagram based on PGL-3-C1 computed with the same parameter values and methods as in Fig. 2. Comparison to experimental data are shown for the dilute branch at 20 °C. Panel to the right (red) is a zoom into the left panel. Also shown are the theoretically predicted tie lines showing a slightly positive slope. **D** Predicted salt-concentration phase diagram based on PGL-3-C1 computed as in **C** but at 10 °C and compared to experimental data for both (dilute and condensed) branches. Note, the graph is shown in semi-logarithmic scale, including the tie lines.

Note that our theoretical model provides a prediction for the slope of the tie lines and the salt content of the protein-rich branch. As an example, for 10 °C and a salt concentration of 150 mM in the protein-poor branch, we obtain a salt concentration of 187 mM in the protein-rich branch. This corresponds to an increase of salt in the condensed phase of 25% and a slope of the tie line equal to 51.9 (M KCl / M PGL-3). We expect to test these results in future experiments.

### FUS-N1 domain is responsible for LLPS

We have shown, in the case of PGL-3, how the prediction of its temperature vs. protein concentration or salt vs. protein concentration phase diagrams can be achieved using only minimal experimental input data. However, the actual predicting power of our approach is revealed for structurally more complex proteins such as FUS. In contrast to PGL-3, which has only one long, continuous IDR, the application of the same machine learning tools yields that FUS has two long IDRs. Note that the structure of FUS is well known (28), but we proceed according to the output of the disorder prediction software, since our stated goal is to apply our method to proteins with unknown structure. Nevertheless, it is known that FUS contains a prion-like low-complexity domain (29) (LC) (aa 1-214), which overlaps with the predicted IDR near the N terminus (aa 1-285, FUS-N1) (Fig. 4A) and is mostly devoid of charged amino acids. We predict an additional IDR near the C terminus (aa 367-526, FUS-C1)(Fig. 4A). Within the latter IDR it is localized a Zn-finger domain, which is reported to have a role in RNA binding and sequence recognition (30) The role of these domains in LLPS of FUS is still not known and in particular the role of the LC domain is still debated, even if it is known to be necessary for phase separation, see the discussion below.

**Figure 4:**
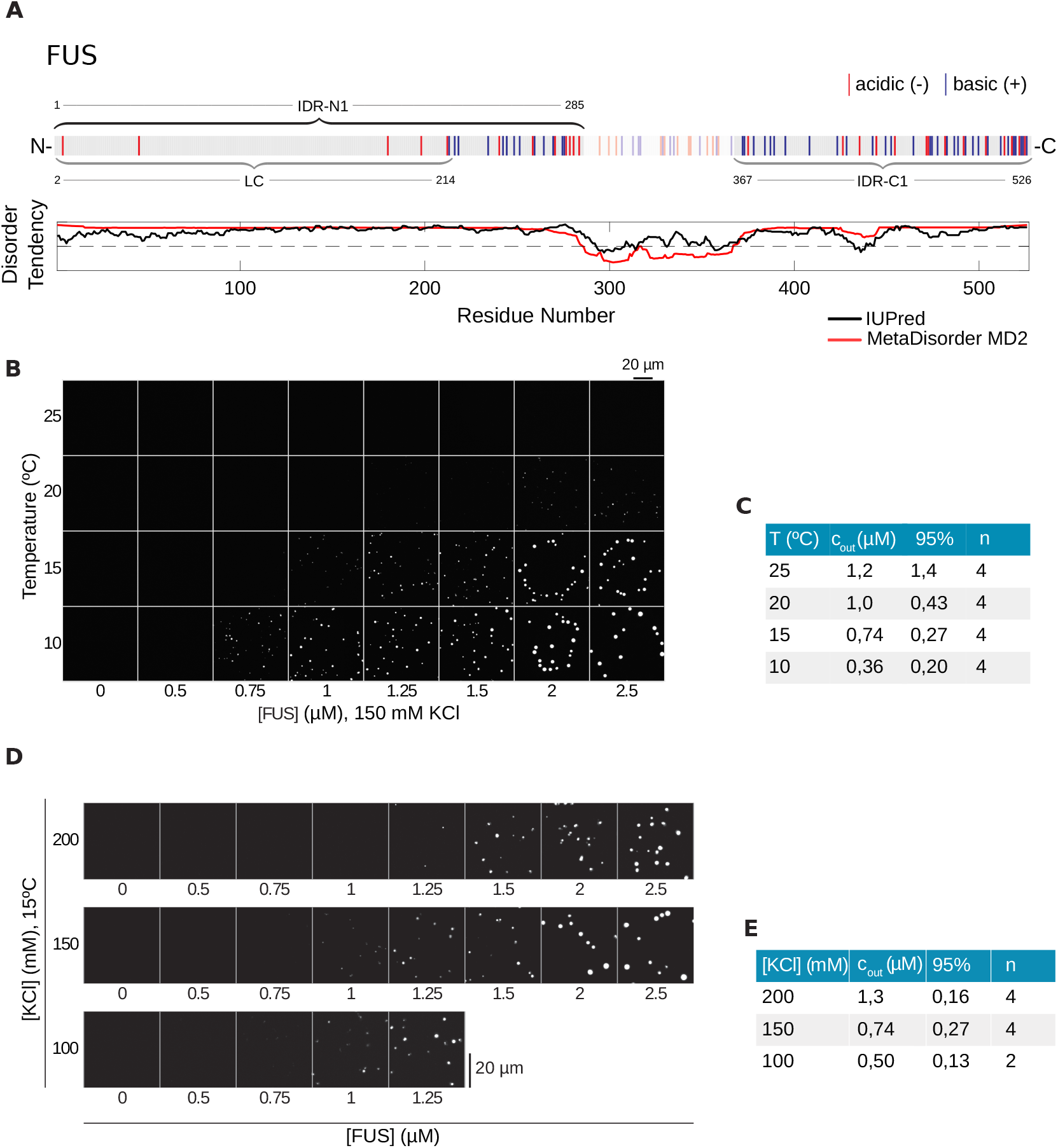
The LLPS of FUS depends weakly on salt concentration **A** Sequence of hFUS with predicted N-terminal IDR (IDR-N1), which contains the well-described low-complexity (LC) region (14), and a C-terminal IDR (IDR-C1). **B** Temperature- and concentration-dependent LLPS of hFUS-mEGFP at 150 mM KCl. **C** Quantification of data in **B** to derive *c*_*out*_ with *n* = 4 repetitions, 95% denotes the corresponding confidence interval. **D** Salt- and concentration-dependent phase separation of hFUS-mEGFP show a weak salt dependence of LLPS. **E** Quantification of data in **D** to obtain *c*_*out*_. Column *n* corresponds to the number of repetitions and 95% denotes the corresponding confidence interval.

Initially, the comparison of the theoretical and experimental results is performed using FUS-N1, which gives the smallest overall error (see Table 1 in Methods and Materials), and later we also explore the consequences of using the other IDRs we have identified. We show that we can not only predict if FUS undergoes LLPS by itself under physiological conditions, but we can now test whether our approach can determine which domain is responsible for LLPS of FUS.

In a similar fashion as for PGL-3, we first investigate experimentally the temperature-protein concentration phase diagram in order to fit the parameters of the model. In contrast to PGL-3, we observe for FUS that the influence of temperature on the saturation concentration *c*_*out*_ is comparable to the change we see for salt, with a 3.4-fold increase from 0.36 to 1.21 *μ*M between 10 and 25 °C (Fig. 4B,C) and a salt dependence in a range of 100-200 mM KCl with a 2.5-fold change from 0.5 to 1.26 *μ*M (Fig. 4D,E). Considering the FUS-N1 region for the model (IDR-N1, highlighted in Fig. 5A), the theoretical results show good agreement with the experimental results (Fig. 5B). The theoretical phase diagram has a maximum near 150°C and has an even more striking *pointy* feature. If we zoom in to the region with a biologically meaningful temperature (Fig. 5A, right panel), we can see how well the overlaid experimental points fall upon the theoretical dilute branch. In this case, the experimental point at 10°C also falls directly upon the condensed branch, thus giving an even better fit that PGL-3, which is manifest in the value of the *χ*^2^ parameter (see Table 1 in Methods and Materials).

**Figure 5:**
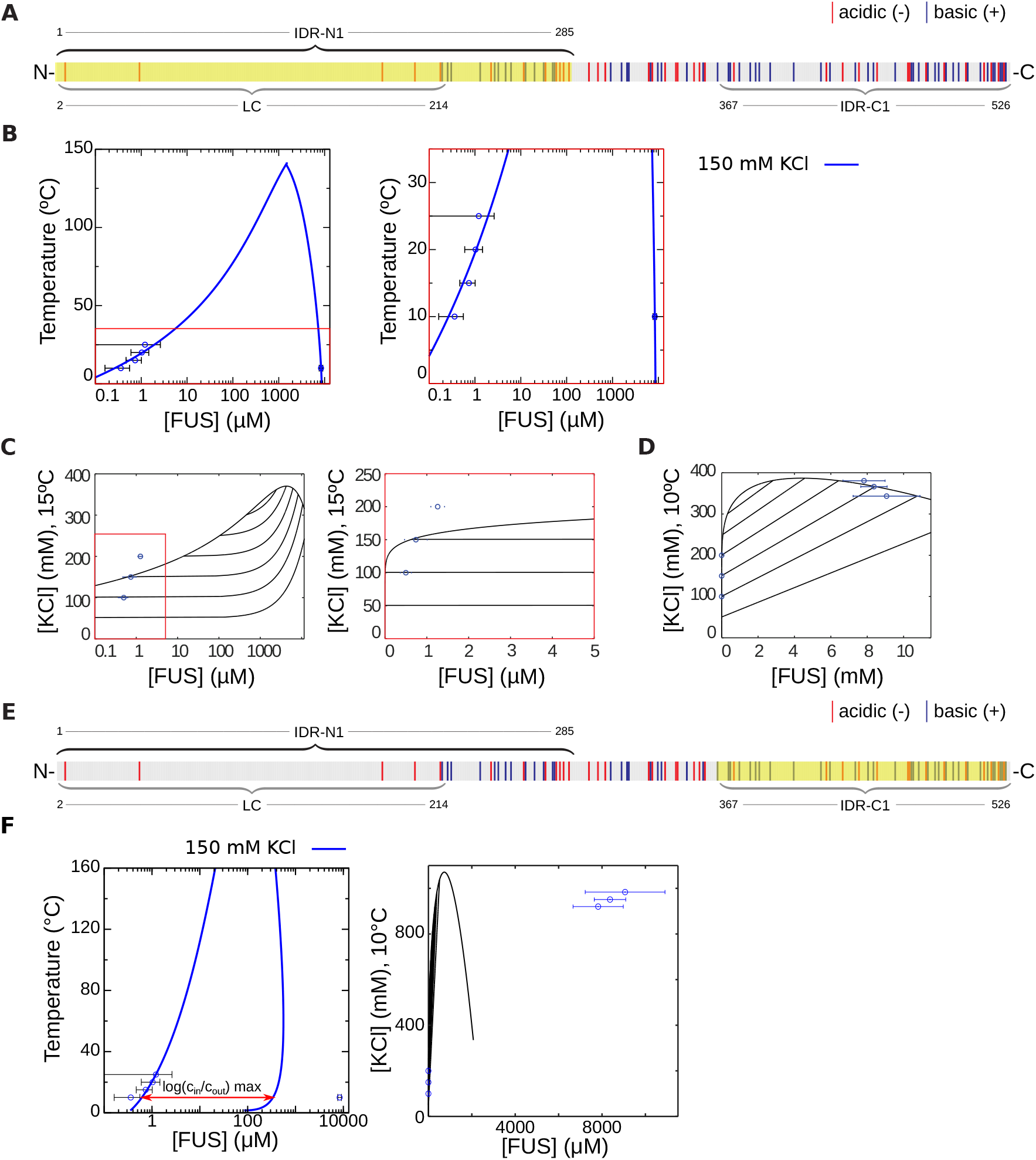
The domain FUS-N1 is responsible for LLPS in FUS. **A** Intrinsically disordered region FUS-N1 (IDR-N1, highlighted) **B** Predicted temperature-concentration phase diagram based on RPA analysis using FUS-N1 with parameter values *ε*_*r*_ = 8.76 and *β*_2_ = 207 mM, for 150 mM KCl. **C** Predicted salt-concentration phase diagram for *T* = 20°C obtained using the same parameter values as in **B**. The right panel is a zoom to the red square in the left panel. Experimental data is shown for the dilute branch **D** Predicted salt-concentration phase diagram for *T* = 20°C, same parameters as **B**. Experimental data is shown for the dilute and condensed branches. **E** Intrinsically disordered region FUS-C1 (IDR-C1, highlighted) **F** Phase diagrams in the FUS-C1 case. Left panel: theoretical temperature-concentration diagram at 150 mM KCl obtained with parameter values *ε*_*r*_ = 7.62 and *β*_2_ = 10.1 mM. Right panel: salt-concentration diagram obtained with the same parameters. Notice that the experimental points cannot be fitted to the condensed branch of the model, since the ratio of *c*_*in*_ to *c*_*out*_ is much greater than its maximum value (see discussion in the main text).

As in the PGL-3 case, the experimentally fitted parameter values obtained for the temperature-concentration phase diagram are also used to generate the salt-concentration phase diagram for comparison to the experimental data. The overlay of experimental points and the theory captures the trend of the change of concentration with the salinity in the dilute branch (Fig. 5C). Again the discrepancy in the values of the concentration stems from the fact that we use the value of the permittivity at 150 mM KCl but we know that its value depends strongly on salt concentration. In the condensed branch we obtain a a very good agreement between theory an experiments (Fig. 5D), probably on account of the smaller variation of the salinity. For 10°C we can compute the slope of the tie line for a salt concentration of 150 mM in the dilute branch, giving 25.6 (M KCl / M FUS), and a predicted salt concentration of 366 mM, and thus an increase of 144% in the condensed branch, which we also expect to be tested experimentally in the future.

We also investigated the FUS-LC domain and found that the theoretical curve cannot be fitted to allow for comparison with the experimental results, since the scaling factor involves a permittivity smaller than 1 as shown in Table 1 in Methods and Materials. Thus, interestingly, the LC domain should not play a dominant role in LLPS of FUS, according to our approach. Other than FUS-N1, a further IDR should be considered according to the results from MetaDisorder, FUSC1 (highlighted in Fig. 5E). In principle one could compare the fits and reason in terms of the goodness of fit (i.e. the *χ*^2^ parameter given in the Methods and Materials section) to decide which region is most likely responsible. But a side by side comparison of the overlay of the experimental data points for FUS-N1 and FUS-C1 shows a much more clear-cut result, which is that the experimental results cannot be fitted to the FUS-C1 curve (See Fig. 5F). This is a consequence of salt partitioning and our parsimonious fitting approach.

Salt partitioning and the requisite that the dilute branch has a constant salt concentration give a very constrained form of the phase diagram (Fig. 5F, left panel). This is in contrast with constant-salt models, which give a binodal curve that can be conveniently scaled to fit most experimental data. In particular, salt partitioning imposes a maximum ratio of the protein concentration in the condensed branch with the concentration in the dilute branch. This maximum ratio is marked explicitly with an arrow in Fig. 5F, left panel. The arrow has a length of log (*c*_*in*_ /*c*_*out*_), which is the log of the maximum of the ratio *c*_*in*_/*c*_*out*_. If the experimental *c*_*in*_/*c*_*out*_ is greater than this maximum value, the theoretical curve cannot be scaled to fit the experiments. This means that either the condensed or the dilute branch can be fitted, but not both. In Fig. 5F we observe that the dilute branch is well captured by the theoretical model, but there is a large discrepancy with the condensed branch, which is even more manifest in a linear concentration scale in the salt-protein phase diagram (Fig. 5F, right panel).

This surprising result is in clear contrast with models where the salinity is considered constant in both phases (as shown in Fig. S1) and thus clearly shows the predictive power of our model, discarding regions for being responsible for LLPS while giving a testable prediction for the salinity of the condensed phase.

## DISCUSSION

In this study we investigated the disordered regions of the *C. elegans* protein PGL-3 and the human fused in sarcoma (FUS) protein regarding their role in LLPS using random phase approximation (RPA) for the electrostatic interactions and including variable salt concentrations. By direct comparison of the resulting theoretical predictions of the phase behaviour with *in vitro* experimental results under physiological salt conditions we show that the model is capable to identify specific domains that trigger LLPS.

For FUS, the role in phase separation of the different domains, such as the LC domain, has been controversial, starting with its structural identification. In fact, the LC domain was first identified in a bioinformatics survey for prion-like domains in different proteins (31, 32), identifying the LC region to amino acids 1 to 239. Kato et al. (29) identified this region using SEG Wootton and Federhen (33) to be amino acids 2 to 214. We have adopted their definition, as it has become a standard in the subsequent literature, but it has not been possible to reproduce their result using SEG. Kato et al. (29) identified the FUS-LC region as responsible for the protein hydrogel formation by performing experiments with the excised domain, but used a very large concentration (≈2.3 mM). Later, Patel et al. (16) (cf. Murthy et al. (34)) proved that the LC region is necessary for phase separation, since the protein with that region excised would not phase-separate. Interestingly, they also found that a single mutation in the LC domain does not change the phase diagram (even if it has a strong effect on the kinetics of the transformation, e.g. changing the timescale of fibril formation). In any case, the concentrations at which phase separation occurs are much higher for FUS-LC than for FUS(35).

Our results on FUS clearly single out the IDR N1 at the N-terminus (1-285) to lead to phase separation of the full protein in comparison with experiments, while other candidate regions such as the LC or IDR at the C-terminus either require a permittivity smaller than one or can not be fitted to the experimental data at all. These results are in disagreement with Kato et al. (29) but Wang et al. (14) and Luo et al. (36) support our findings. They showed that the LC domain needs indeed very high concentrations in order to phase-separate on its own, which was the regime tested by Kato et al. (29). Wang et al. (14) further ascertain that it is the interaction of the LC region and the RNA binding domain which makes the phase transition possible at low protein concentrations. In other recent studies (37, 38), coarse grained models in combination with MD simulations have been used. In particular in Dignon et al. (38) it is shown that the longer a sequence containing the LC is, the higher is the propensity towards phase separation. In accordance with our results, Kang et al. (39) demonstrate experimentally that an extended LC domain containing the RGG region next to it phase separated at a much smaller saturation concentration than the LC domain. The domain reported by Kang et al. (39) is very similar to our N1 domain, thus supporting our theoretical results. Moreover, our results show that the phase diagram corresponding to the FUS-C1 region cannot be fitted to the experimental results, thus showing that our model is able to discard IDRs as responsible for phase separation in different ways. These findings suggest further experimental studies on the role of IDR N1, specifically on the role of the amino acids that are lacking in the LC region. It would also require to systematically explore the impact of variations as well as mutations of these regions in our future theoretical studies.

We further note that the connection of the domain structure of proteins with their phase behaviour may be rooted in the underlying model system for polyampholytes, that we used in our analysis. Indeed, Lytle et al. (40) investigated the impact of the blockyness of polyampholytes on their phase behaviour, and Das et al. (41) compared results from Monte Carlo simulations to RPA for specific polyampholyte sequences showing that patterns of larger blocks of charge lead to significantly higher tendency to phase separate.

Also, one can argue that the failure of FUS-LC to phase separate in our framework is directly related to the fact that our approach only considers elctrostatic interactions between charged residues and not other kind of interactions. Then, since the FUS-LC domain is mostly devoid of charges it is not surprising that it does not phase separate. We see this as a confirmation of the relevance of charged residues in the case of FUS, and thus our simple RPA approach is appropriate to understand the main mechanism of LLPS in FUS. We do not claim, however, that we capture the complete picture of LLPS in FUS, since, for instance, FUS-LC is known to phase-separate, even if it does so at a very high concentrations due to other interaction mechanisms(42), and the RGG domain near the C terminus is known to influence LLPS(34). We have also shown the impact of salt distribution on the resulting phase diagram and obtained salt partitioning into the condensed phase for both proteins, making manifest the strong salt dependence of the permittivity. The inclusion of salt as a further variable in the model led to phase diagrams with a characteristic *pointy* feature, corresponding to a discontinuity in the slope of the binodal curve. We showed that this is a generic feature and is rooted in the fact that salt concentration is not imposed to be constant everywhere, but is allowed to vary in the condensed phase. In previous studies this fact has often been neglected but, in fact, it is thermodynamically not consistent to do so.

The experimental corroboration of our predictions on the salt concentration in the condensed phase will be part for our future studies, requiring a detailed discussion of the theoretical model for polyampholyte solutions that we used as a model system for proteins. Here, we remark that the question of salt partitioning during LLPS is also not completely understood, even for polyelectrolytes and polyampholytes. This is the case in the context of Random-Phase-Approximation, Liquid-State-Theory, Monte Carlo simulation and Voorn-Overbeek theory, as well as in experimental studies (43–49). Voorn-Overbeek theory predicts an excess of salt in the condensed phase, corresponding to a positive slope of the tie line, while more recent theoretical and experimental studies find that excluded volume effects are responsible for expelling the salt counterions into the dilute phase, thus predicting a negative slope of the tie lines. However, for low overall salt concentration this can be reversed, see for example Li et al. (49). Also, molecular dynamics simulations seem to indicate that electrical neutrality and a lack of preferential interactions between salt ions and interactions are enough to obtain a reasonable prediction of the concentration of salt ions in the condensed phase (50). In summary, the slope of the tie lines and salt partitioning depend strongly on the model used, and more experimental results are needed to guide the theoretical efforts.

While RPA is the appropriate tool to address the structural properties of IDPs Rumyantsev et al. (51), recent discussion in Das et al. (41), where explicit chain simulation and RPA are compared for a number of polyampholyte sequences, suggest higher order contributions of the functional integral of the partition function in order to address the accuracy of RPA (Lin et al. (11), Borue and Erukhimovich (23), Shen and Wang (52)), specifically in the protein-poor phase.

We also note that further physical interactions also play a role and are still being discovered Berry et al. (53). Currently, our model does not account for non-specific contacts between positively charged arginine-glycine-glycine (RGG) domains, such as those found in FUS or PGL-3, and negatively charged RNA, which can strengthen the binding affinity of existing RNA binding domains and could provide alternative interaction modes. Also, heterotypic interactions with other regions of the same polypeptide or other proteins are known to drive phase separation Wang et al. (14) but can be included in principle into our framework.

## AUTHOR CONTRIBUTIONS

E.M. designed the theoretical research, carried out all simulations and parameter estimations and helped prepare the manuscript. A.W.F. and J.M.I. carried out the experiments, analyzed the data and helped prepare the manuscript. S.R. helped conceive the research and to prepare the manuscript. B.W. conceived the research, supervised the project and prepared the manuscript.

## ACKNOWLEDGMENTS

We acknowledge Anthony Hyman for providing all the support needed for the experimental part of this project. E.M. would like thank the Weierstrass Institute for hosting a research visit. E.M. and B.W. would like to thank Andreas Münch for helpful discussions on the theoretical aspects of the research project. For discussions and help with the experimental part of the project we thank, Martine Ruer, Patrick McCall, Tylor Harmon, Jie Wang, and Titius Franzmann. We thank the light microscopy, chromatography, and protein purification facilities at MPI-CBG for their support. We thank Olympus for providing the CSU-W1 SoRa spinning-disc system based on an IXplore IX83 microscope.

A.W.F. was supported by the ELBE postdoctoral fellows program and the Max Planck Research Network for Synthetic Biology (MaxSynBio) consortium, jointly funded by the Federal Ministry of Education and Research of Germany and the Max Planck Society.

## REFERENCES

1. Banani, S. F., H. O. Lee, A. A. Hyman, and M. K. Rosen, 2017. Biomolecular condensates: organizers of cellular biochemistry. Nature reviews Molecular cell biology 18:285–298.

2. Shin, Y., and C. P. Brangwynne, 2017. Liquid phase condensation in cell physiology and disease. Science 357:eaaf4382.

3. Alberti, S., and A. A. Hyman, 2021. Biomolecular condensates at the nexus of cellular stress, protein aggregation disease and ageing. Nat Rev Mol Cell Biol 22:196–213.

4. Brangwynne, C. P., P. Tompa, and R. V. Pappu, 2015. Polymer physics of intracellular phase transitions. Nature Physics 11:899–904.

5. Rauscher, S., and R. Pomès, 2017. The liquid structure of elastin. Elife 6:e26526.

6. Flory, P. J., 1942. Thermodynamics of high polymer solutions. The Journal of chemical physics 10:51–61.

7. Huggins, M. L., 1942. Some properties of solutions of long-chain compounds. The Journal of Physical Chemistry 46:151–158.

8. Overbeek, J. T. G., and M. Voorn, 1957. Phase separation in polyelectrolyte solutions. Theory of complex coacervation. Journal of Cellular and Comparative Physiology 49:7–26.

9. Zhang, X., M. Vigers, J. McCarty, J. N. Rauch, G. H. Fredrickson, M. Z. Wilson, J.-E. Shea, S. Han, and K. S. Kosik, 2020. The proline-rich domain promotes Tau liquid–liquid phase separation in cells. Journal of Cell Biology 219.

10. Lin, Y.-H., J. D. Forman-Kay, and H. S. Chan, 2016. Sequence-specific polyampholyte phase separation in membraneless organelles. Physical review letters 117:178101.

11. Lin, Y.-H., J. P. Brady, H. S. Chan, and K. Ghosh, 2020. A unified analytical theory of heteropolymers for sequence-specific phase behaviors of polyelectrolytes and polyam-pholytes. The Journal of chemical physics 152:045102.

12. Dinic, J., A. B. Marciel, and M. V. Tirrell, 2021. Polyampholyte Physics: Liquid-Liquid Phase Separation and Biological Condensates. Current Opinion in Colloid & Interface Science 101457.

13. Brangwynne, C. P., C. R. Eckmann, D. S. Courson, A. Rybarska, C. Hoege, J. Gharakhani, F. Jülicher, and A. A. Hyman, 2009. Germline P Granules Are Liquid Droplets That Localize by Controlled Dissolution/Condensation. Science 324:1729–1732.

14. Wang, J., J.-M. Choi, A. S. Holehouse, H. O. Lee, X. Zhang, M. Jahnel, S. Maharana, R. Lemaitre, A. Pozniakovsky, D. Drechsel, I. Poser, R. V. Pappu, S. Alberti, and A. A. Hyman, 2018. A Molecular Grammar Governing the Driving Forces for Phase Separation of Prion-like RNA Binding Proteins. Cell 174:688 – 699.e16. http://www.sciencedirect.com/science/article/pii/S0092867418307311.

15. Saha, S., C. A. Weber, M. Nousch, O. Adame-Arana, C. Hoege, M. Y. Hein, E. Osborne-Nishimura, J. Mahamid, M. Jahnel, L. Jawerth, et al., 2016. Polar positioning of phase-separated liquid compartments in cells regulated by an mRNA competition mechanism. Cell 166:1572–1584.

16. Patel, A., H. O. Lee, L. Jawerth, S. Maharana, M. Jahnel, M. Y. Hein, S. Stoynov, J. Mahamid, S. Saha, T. M. Franzmann, et al., 2015. A liquid-to-solid phase transition of the ALS protein FUS accelerated by disease mutation. Cell 162:1066–1077.

17. Kozlowski, L. P., and J. M. Bujnicki, 2012. MetaDisorder: a meta-server for the prediction of intrinsic disorder in proteins. BMC Bioinformatics 13:111. http://dx.doi.org/10.1186/1471-2105-13-111.

18. Dosztányi, Z., 2018. Prediction of protein disorder based on IUPred. Protein Science 27:331–340.

19. Baeurle, S. A., 2002. Method of Gaussian Equivalent Representation: A New Technique for Reducing the Sign Problem of Functional Integral Methods. Physical Review Letters 89:080602.

20. Lin, Y.-H., J. Song, J. D. Forman-Kay, and H. S. Chan, 2017. Random-phase-approximation theory for sequence-dependent, biologically functional liquid-liquid phase separation of intrinsically disordered proteins. Journal of Molecular Liquids 228:176–193.

21. Castelnovo, M., and J.-F. Joanny, 2001. Complexation between oppositely charged polyelectrolytes: Beyond the Random Phase Approximation. The European Physical Journal E 6:377–386. https://doi.org/10.1007/s10189-001-8051-7.

22. Borukhov, I., D. Andelman, and H. Orland, 1998. Random polyelectrolytes and polyampholytes in solution. The European Physical Journal B - Condensed Matter and Complex Systems 5:869–880. https://doi.org/10. 1007/s100510050513.

23. Borue, V. Y., and I. Y. Erukhimovich, 1988. A statistical theory of weakly charged polyelectrolytes: fluctuations, equation of state and microphase separation. Macro-molecules 21:3240–3249.

24. Ermoshkin, A. V., and M. Olvera de la Cruz, 2004. Gelation in strongly charged polyelectrolytes. J. Ploym. Sci. Part B 42:766–776.

25. Golub, G. H., and J. H. Welsch, 1969. Calculation of Gauss quadrature rules. Mathematics of computation 23:221–230.

26. Glaser, A., X. Liu, and V. Rokhlin, 2007. A fast algorithm for the calculation of the roots of special functions. SIAM Journal on Scientific Computing 29:1420–1438.

27. Govaerts, W. J., 2000. Numerical methods for bifurcations of dynamical equilibria, volume 66. Siam.

28. Aulas, A., and C. Vande Velde, 2015. Alterations in stress granule dynamics driven by TDP-43 and FUS: a link to pathological inclusions in ALS? Frontiers in cellular neuroscience 9:423.

29. Kato, M., T. W. Han, S. Xie, K. Shi, X. Du, L. C. Wu, H. Mirzaei, E. J. Goldsmith, J. Longgood, J. Pei, et al., 2012. Cell-free formation of RNA granules: low complexity sequence domains form dynamic fibers within hydrogels. Cell 149:753–767.

30. Loughlin, F. E., P. J. Lukavsky, T. Kazeeva, S. Reber, E.-M. Hock, M. Colombo, C. Von Schroetter, P. Pauli, A. Cléry, O. Mühlemann, et al., 2019. The solution structure of FUS bound to RNA reveals a bipartite mode of RNA recognition with both sequence and shape specificity. Molecular cell 73:490–504.

31. Cushman, M., B. S. Johnson, O. D. King, A. D. Gitler, and J. Shorter, 2010. Prion-like disorders: blurring the divide between transmissibility and infectivity. Journal of Cell Science 123:1191–1201.

32. Alberti, S., R. Halfmann, O. King, A. Kapila, and S. Lindquist, 2009. A Systematic Survey Identifies Prions and Illuminates Sequence Features of Prionogenic Proteins. Cell 137:146–158.

33. Wootton, J. C., and S. Federhen, 1996. [33] Analysis of compositionally biased regions in sequence databases. In Methods in enzymology, Elsevier, volume 266, 554–571.

34. Murthy, A. C., W. S. Tang, N. Jovic, A. M. Janke, D. H. Seo, T. M. Perdikari, J. Mittal, and N. L. Fawzi, 2021. Molecular interactions contributing to FUS SYGQ LC-RGG phase separation and co-partitioning with RNA polymerase II heptads. Nature Structural & Molecular Biology 28:923–935.

35. Burke, K. A., A. M. Janke, C. L. Rhine, and N. L. Fawzi, 2015. Residue-by-residue view of in vitro FUS granules that bind the C-terminal domain of RNA polymerase II. Molecular cell 60:231–241.

36. Luo, F., X. Gui, H. Zhou, J. Gu, Y. Li, X. Liu, M. Zhao, D. Li, X. Li, and C. Liu, 2018. Atomic structures of FUS LC domain segments reveal bases for reversible amyloid fibril formation. Nature structural & molecular biology 25:341–346.

37. Benayad, Z., S. von Bülow, L. S. Stelzl, and G. Hummer, 2021. Simulation of FUS Protein Condensates with an Adapted Coarse-Grained Model. Journal of Chemical Theory and Computation 17:525–537.

38. Dignon, G., W. Zheng, Y. Kim, R. Best, and J. Mittal, 2018. Sequence determinants of protein phase behavior from a coarse-grained model. PLoS Comput Biol 14:e1005941.

39. Kang, J., L. Lim, Y. Lu, and J. Song, 2019. A unified mechanism for LLPS of ALS/FTLD-causing FUS as well as its modulation by ATP and oligonucleic acids. PLoS biology 17:e3000327.

40. Lytle, T. K., L.-W. Chang, N. Markiewicz, S. L. Perry, and C. E. Sing, 2019. Designing Electrostatic Interactions via Polyelectrolyte Monomer Sequence. ACS Central Science 5:709–718.

41. Das, S., A. Eisen, Y.-H. Lin, and H. S. Chan, 2018. A Lattice Model of Charge-Pattern-Dependent Polyampholyte Phase Separation. The Journal of Physical Chemistry B 122:5418–5431.

42. Murthy, A. C., G. L. Dignon, Y. Kan, G. H. Zerze, S. H. Parekh, J. Mittal, and N. L. Fawzi, 2019. Molecular interactions underlying liquidliquid phase separation of the FUS low-complexity domain. Nature structural & molecular biology 26:637–648.

43. Perry, S. L., and C. E. Sing, 2015. PRISM-Based Theory of Complex Coacervation: Excluded Volume versus Chain Correlation. Macromolecules 48:5040–5053.

44. Radhakrishna, M., K. Basu, Y. Liu, R. Shamsi, S. L. Perry, and C. E. Sing, 2017. Molecular Connectivity and Correlation Effects on Polymer Coacervation. Macromolecules 50:3030–3037.

45. Zhang, P., K. Shen, N. M. Alsaifi, and Z.-G. Wang, 2018. Salt Partitioning in Complex Coacervation of Symmetric Polyelectrolytes. Macromolecules 51:5586–5593.

46. Madinya, J. J., L.-W. Chang, S. L. Perry, and C. E. Sing, 2019. Sequence-dependent self-coacervation in high charge-density polyampholytes. Mol. Syst. Des. Eng. –. http://dx.doi.org/10.1039/C9ME00074G.

47. Lytle, T. K., and C. E. Sing, 2017. Transfer matrix theory of polymer complex coacervation. Soft Matter 13:7001–7012.

48. Shen, K., and Z.-G. Wang, 2018. Polyelectrolyte Chain Structure and Solution Phase Behavior. Macromolecules 51:1706–1717.

49. Li, L., S. Srivastava, M. Andreev, A. B. Marciel, J. J. de Pablo, and M. V. Tirrell, 2018. Phase Behavior and Salt Partitioning in Polyelectrolyte Complex Coacervates. Macromolecules 51:2988–2995.

50. Zheng, W., G. L. Dignon, N. Jovic, X. Xu, R. M. Regy, N. L. Fawzi, Y. C. Kim, R. B. Best, and J. Mittal, 2020. Molecular details of protein condensates probed by microsecond long atomistic simulations. The Journal of Physical Chemistry B 124:11671–11679.

51. Rumyantsev, A. M., N. E. Jackson, B. Yu, J. M. Ting, W. Chen, M. V. Tirrell, and J. J. de Pablo, 2019. Controlling Complex Coacervation via Random Polyelectrolyte Sequences. ACS Macro Letters 8:1296–1302.

52. Shen, K., and Z.-G. Wang, 2017. Electrostatic correlations and the polyelectrolyte self energy. The Journal of Chemical Physics 146:084901.

53. Berry, J., C. Brangwynne, and M. P. Haataja, 2018. Physical Principles of Intracellular Organization via Active and Passive Phase Transitions. Reports on Progress in Physics.

